# EpiGraphDB: A database and data mining platform for health data science

**DOI:** 10.1101/2020.08.01.230193

**Authors:** Yi Liu, Benjamin Elsworth, Pau Erola, Valeriia Haberland, Gibran Hemani, Matt Lyon, Jie Zheng, Tom R Gaunt

## Abstract

**Motivation:** The wealth of data resources on human phenotypes, risk factors, molecular traits and therapeutic interventions presents new opportunities for population health sciences. These opportunities are paralleled by a growing need for data integration, curation and mining to increase research efficiency, reduce mis-inference and ensure reproducible research.

**Results:** We developed EpiGraphDB (https://epigraphdb.org/), a graph database containing an array of different biomedical and epidemiological relationships and an analytical platform to support their use in human population health data science. In addition, we present three case studies that illustrate the value of this platform. The first uses EpiGraphDB to evaluate potential pleiotropic relationships, addressing mis-inference in systematic causal analysis. In the second case study we illustrate how protein-protein interaction data offer opportunities to identify new drug targets. The final case study integrates causal inference using Mendelian randomization with relationships mined from the biomedical literature to “triangulate” evidence from different sources.

**Availability:** The EpiGraphDB platform is openly available at https://epigraphdb.org. Code for replicating case study results is available at https://github.com/MRCIEU/epigraphdb as Jupyter notebooks using the API, and https://mrcieu.github.io/epigraphdb-r using the R package.

## 1 Introduction

The wealth and diversity of population data now available to epidemiologists is enabling new discoveries and methods development in population health data science. However, harmonisation and integration of data presents a challenge to researchers aiming to “triangulate” evidence from different sources or uncover potential mechanistic pathways. This challenge can be tackled through the development of data integration platforms which curate and combine data sources to enable integrative analyses. One area in which data integration offers potential value is causal inference. Over the last two decades Mendelian randomization (MR) (Davey Smith and Ebrahim, 2003) has risen to prominence as a key causal inference method. MR exploits genetic variants as causal “anchors” (randomly allocated and invariant from conception) to estimate causal effects between an “exposure” (risk factor) influenced by the genetic variant(s) and a health outcome. The approach has various assumptions, of which a key constraint is that the genetic variants should not pleiotropically affect the health outcome through a pathway other than the risk factor in question. The two-sample MR approach enables MR to be performed in situations where a risk factor (exposure) and an outcome are analysed for genetic association in separate studies (Pierce and Burgess, 2013), enabling the thousands of published genome-wide association study (GWAS) datasets (MacArthur *et al.*, 2016) to be leveraged for causal inference.

Database resources such as the IEU OpenGWAS database (*https://gwas.mrcieu.ac.uk*) (Elsworth, Lyon, *et al.*, 2020), and the linked MR-Base analytical platform (Hemani *et al.*, 2018) now enable systematic MR using approaches such as the MR Mixture of Experts (Hemani *et al.*, 2017). Such systematic MR analyses offer the potential to take a “systems” approach to the evaluation of potential intervention targets, as we have recently demonstrated with the plasma proteome (Zheng, Haberland, *et al.*, 2019). However, such systematic approaches raise new challenges in interpretation of the wealth of causal estimates generated. The integration of causal estimates with data from other sources is one way to tackle such challenges. Combining evidence with different biases (such as MR estimates, observational correlations and literature-mined experimental results) can provide more robust causal interpretation in an approach described as “triangulation” (Lawlor *et al.*, 2017). Agreement between sources strengthens the case for causality, whilst disagreement helps identify sources of bias.

Integration of data also offers the scope to gain more mechanistic insight into complex networks of association. Linking phenotypic data with genetic variants and molecular pathway data may make it easier to identify potential intervention targets once a causal relationship has been established. Similarly, an extensive network of associations provides the opportunity to identify drug repositioning opportunities and on-target side effects for pharmaceutical targets.

Here we describe EpiGraphDB (https://epigraphdb.org/), a database and analytical platform, that integrates trait relationships (causal, observational or genetic), literature-mined relationships, biological pathways, protein-protein interactions, drug-target relationships and other data sources to support data mining of risk factor/disease relationships. In the following sections, we describe the EpiGraphDB platform and its epidemiological resources, and then illustrate some potential applications of this platform through specific case studies.

## 2 Implementation

### 3.1 The EpiGraphDB platform

EpiGraphDB integrates data from a range of bioinformatic and epidemiological sources. The database can be accessed using a variety of methods aimed at different needs: programmatically through a custom application programming interface (API) web service, a user-friendly web user interface (UI), an R package, or by directly querying the database via the Neo4j Cypher query language. Figure 1 shows the main components of the EpiGraphDB platform and the overall architecture.

**Fig. 1.**
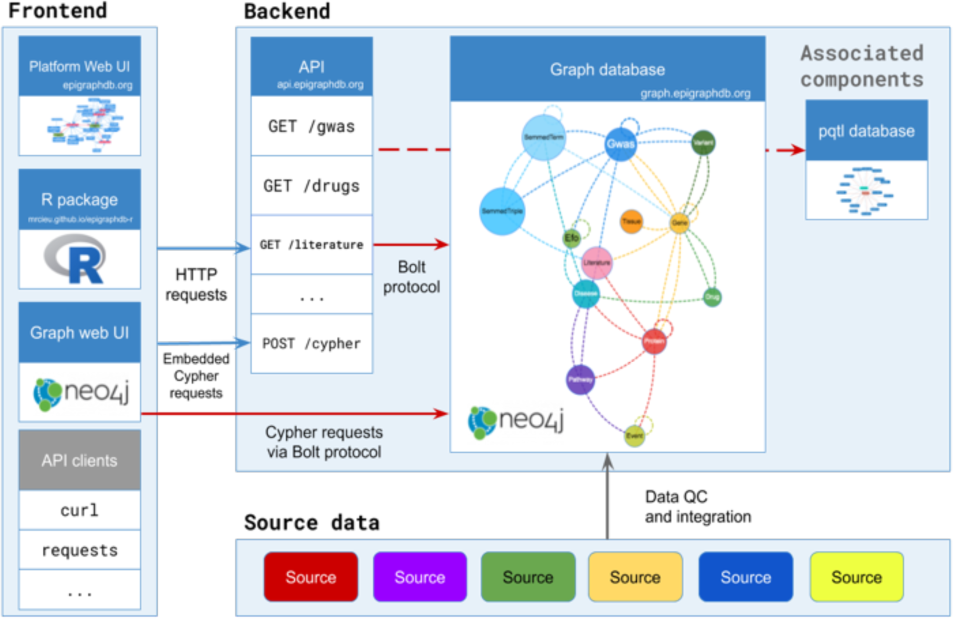
Architecture of the EpiGraphDB platform. Source datasets are integrated into a graph database using Neo4j that uses Neo4j Cypher as query language (which can also be queried by end users). Standard HTTP queries are processed through a RESTful API service which can be called from any REST API client, including our R package epigraphdb. The web UI showcases main topics of the epidemiological evidence in EpiGraphDB and demonstrates the example API queries to get the underlying data.

#### EpiGraphDB Graph

EpiGraphDB is implemented using the Neo4j graph database platform. The graph database paradigm supports interpretable representation of biomedical information by storing data as relationships (e.g. associations, causal estimates, mappings) between entities (e.g. genes, proteins, diseases, genetic variants). The use of Neo4j and the associated Cypher query language also enables more natural representation of hypotheses as queries, in comparison to a relational database architecture using structured query language (SQL). For example, in Neo4j a hypothetical query for the causal effect of a risk factor on disease could be represented in the Cypher query language as `(r:RiskFactor)-[c:CausalEffect]-(d:Disease)`, which illustrates a sub-graph comprising a risk factor node, a disease node and a causal relationship between them. While we recommend most users interface with the EpiGraphDB API regular endpoints or R package for ease of use, we provide a `POST /cypher` endpoint in the EpiGraphDB API^*1*^ (and an interface in the Web UI^*2*^) where users can directly query the Neo4j database using the Cypher query language.

#### EpiGraphDB API

The API provides direct access to pre-defined queries within EpiGraphDB (e.g. MR evidence between phenotypic traits via `GET /mr`, or literature evidence relating to phenotypes via *GET /literature/gwas*), supporting user-specified parameters to select and filter the data. The EpiGraphDB API is a RESTful API service with which users can programmatically retrieve data in commonly used HTTP clients or libraries (e.g. cURL, Requests in Python, httr in R, Postman, etc.) via methods such as `HTTP GET` and `HTTP POST`, without the user having to be proficient in Neo4j Cypher queries. At the same time, for most data queries the API provides the underlying Cypher query to enable users to extend the query to better suit their use cases. This flexible approach enables users to fully utilise rich resources of integrated evidence provided in EpiGraphDB. Online documentation is provided as part of the API (*https://api.epigraphdb.org/* via a Swagger UI) to enable users to explore the API functionality and test their queries.

#### EpiGraphDB Web UI

The Web UI provides a selection of exemplar queries from the API and supports interactive visualisations of sub-graphs generated using those queries. For example, the confounder topic view *https://epigraphdb.org/confounder* demonstrates the use of EpiGraphDB in investigating the potential confounders, mediators and colliders between exposures and outcomes. In addition to viewing the returned data from EpiGraphDB in tabular format and its visualisation in network diagrams, user can also use the “Query” tab to see the underlying API call and Cypher query to assist their further use of EpiGraphDB using the API or Cypher interfaces. In addition, users can use the Explore views *https://epigraphdb.org/explore* to browse and search EpiGraphDB and visit the Gallery *https://epigraphdb.org/gallery* for exemplar use cases.

#### R package *epigraphdb*

This R package provides convenient programmatic access to the major functionalities of the API, enabling users to incorporate EpiGraphDB directly into their analytical pipelines in R for ease of use. Access to API endpoints are wrapped inside R functions and data is returned by default as R tidyverse tibble dataframes, saving users the additional complexity of handling web requests or parsing JSON format response data. Nevertheless, users are able to use the R package as a fully featured API client to EpiGraphDB by sending REST requests to the API, retrieving data in JSON data structures (represented as lists in R) if they wish. They can also use the `POST /cypher` endpoint with the `query_epigraphdb` function if they wish to perform queries not directly represented as R functions.

### 3.2 Integration of epidemiological evidence

EpiGraphDB contains data from a range of biomedical sources, with these data represented as nodes and relationships^*3*^ in a graph database. The relationships broadly represent: epidemiological relationships (e.g. genetic correlations, genetic associations, phenotypic associations and causal estimates from MR) between phenotypes, mappings (e.g. mapping of genetic variants to genes, genes to protein and pathways, genes to drug targets, protein-protein interactions) and relationships derived from the biomedical literature. ***Table 1*** reports a summary of the epidemiological evidence available in EpiGraphDB.

**Table 1.**
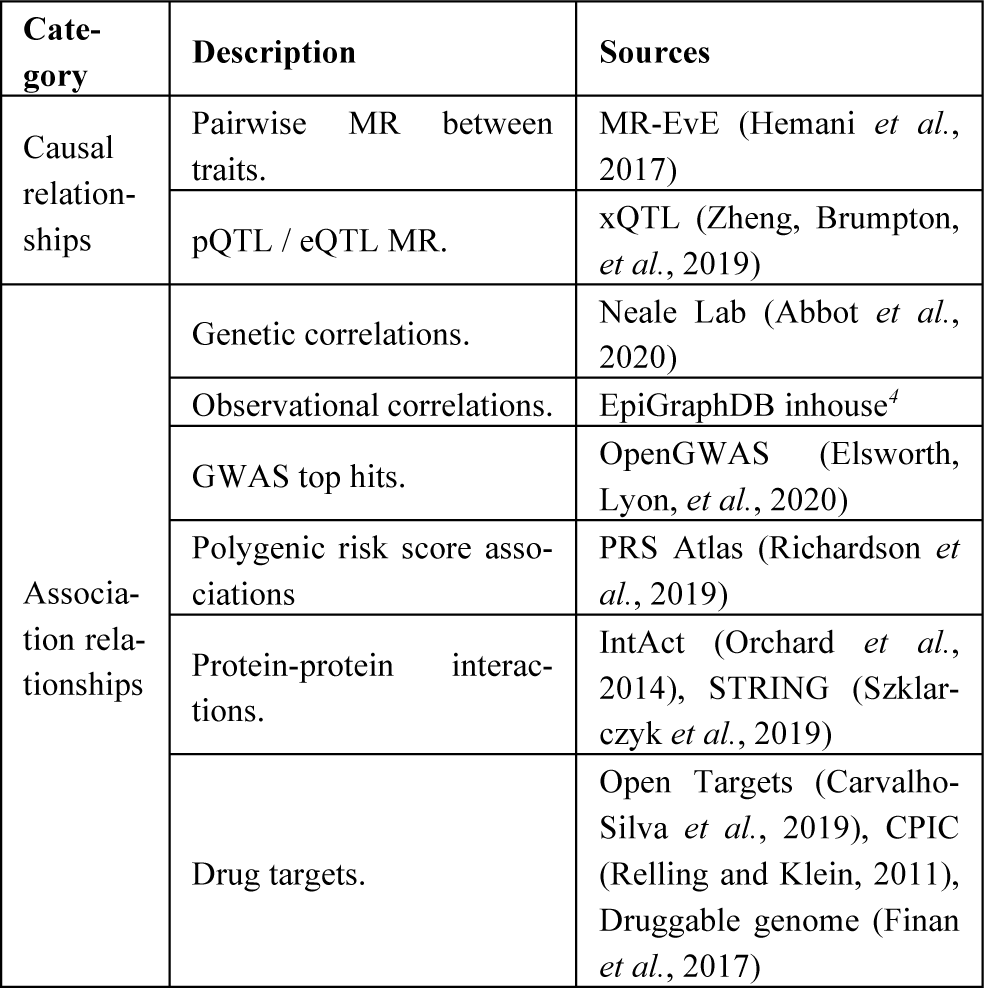
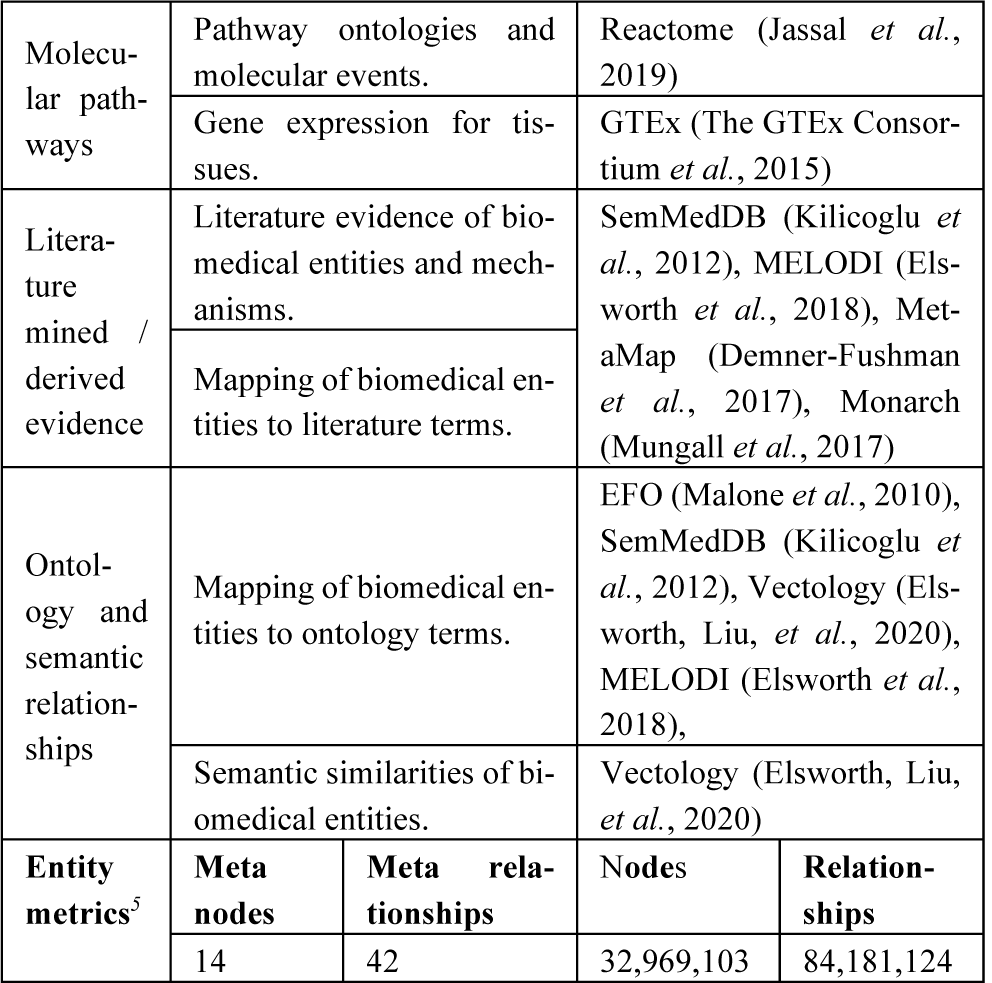
Summary of epidemiological evidence in EpiGraphDB. Detailed discussion on data integration and how these biomedical entities and associations are represented in EpiGraphDB are available in the supplementary materials (Appendices 1 and 2).

| Category | Description | Sources |
| --- | --- | --- |
| Causal relationships | Pairwise MR between traits. | MR-EvE (Hemani *et al.*, 2017) |
|  | pQTL / eQTL MR. | xQTL (Zheng, Brumpton, *et al.*, 2019) |
| Association relationships | Genetic correlations. | Neale Lab (Abbot *et al.*, 2020) |
|  | Observational correlations. | EpiGraphDB inhouse<sup>4</sup> |
|  | GWAS top hits. | OpenGWAS (Elsworth, Lyon, *et al.*, 2020) |
|  | Polygenic risk score associations | PRS Atlas (Richardson *et al.*, 2019) |
|  | Protein-protein interactions. | IntAct (Orchard *et al.*, 2014), STRING (Szklarczyk *et al.*, 2019) |
|  | Drug targets. | Open Targets (Carvalho-Silva *et al.*, 2019), CPIC (Relling and Klein, 2011), Druggable genome (Finan *et al.*, 2017) |

<sup>3</sup> We refer to a type of biomedical entity as a *meta node* (e.g. (*Gwas* in Cypher notation) and a type of association as a *meta relationship* (e.g. [*MR*]), whereas a specific entity is referred to as a *node* (e.g. (*Gwas* {*id*: "ieu-a-2", *trait*: "Body mass index"})) and a specific association as a *relationship* (e.g. (*Gwas* {*trait*: "Body mass

|  |  |  |  |  |
| --- | --- | --- | --- | --- |
| Molecular pathways | Pathway ontologies and molecular events. |  | Reactome (Jassal *et al.*, 2019) |  |
|  | Gene expression for tissues. |  | GTEx (The GTEx Consortium *et al.*, 2015) |  |
| Literature mined / derived evidence | Literature evidence of biomedical entities and mechanisms. |  | SemMedDB (Kilicoglu *et al.*, 2012), MELODI (Elsworth *et al.*, 2018), MetaMap (Demner-Fushman *et al.*, 2017), Monarch (Mungall *et al.*, 2017) |  |
|  | Mapping of biomedical entities to literature terms. |  |  |  |
| Ontology and semantic relationships | Mapping of biomedical entities to ontology terms. |  | EFO (Malone *et al.*, 2010), SemMedDB (Kilicoglu *et al.*, 2012), Vectology (Elsworth, Liu, *et al.*, 2020), MELODI (Elsworth *et al.*, 2018), |  |
|  | Semantic similarities of biomedical entities. |  | Vectology (Elsworth, Liu, *et al.*, 2020) |  |
| Entity metrics<sup>5</sup> | Meta nodes | Meta relationships | Nodes | Relationships |
|  | 14 | 42 | 32,969,103 | 84,181,124 |

We combined data from over 20 independent sources. All datasets required some level of processing in preparation for loading into EpiGraphDB. Detailed information including sources of data, processing steps and data ingest method are described in the supplementary materials (Appendix 2).

## 3 Case studies

The data integrated within EpiGraphDB offer a wide array of potential opportunities for data mining and analysis. Here we present three case studies which illustrate some of the potential for new knowledge discovery using EpiGraphDB. These do not, however, represent the full extent of the data or potential of the platform, which is provided as an open resource for the reader to use for their own novel research investigations.

In case study 1 we explore the potential of pathway data to characterise pleiotropy of genetic instruments used to generate causal estimates of the effect of protein levels on disease outcomes. Case study 2 seeks to identify alternative drug targets using protein-protein interaction data in conjunction with causal estimates of protein levels on disease outcomes as well as literature mined / derived evidence. Case study 3 uses knowledge extracted from the scientific literature to identify potential mechanistic pathways linking causal risk factors to diseases. We discuss the general steps to replicate these case studies in Appendix 5 in the supplementary materials and users are encouraged to use the Jupyter notebooks and R package to replicate and modify the analyses.

### 3.1 Distinguishing vertical and horizontal pleiotropy for SNP-protein associations

A key MR assumption is that the genetic variant (e.g. SNP) is only related to the outcome of interest through the exposure under study (the “exclusion restriction” assumption). This assumption is potentially violated under horizontal pleiotropy, where a SNP is associated with multiple phenotypes (e.g. proteins) independently of the exposure of interest. In contrast, vertical pleiotropy, where a SNP is associated with multiple phenotypes on the causal pathway to the outcome, does not violate the “exclusion restriction criterion” of MR (Figure 2 A). For molecular phenotypes, where the number of genetic instruments is typically limited, it is almost impossible to distinguish vertical and horizontal pleiotropy using established statistical approaches (van Kippersluis and Rietveld, 2018).

**Fig. 2.**
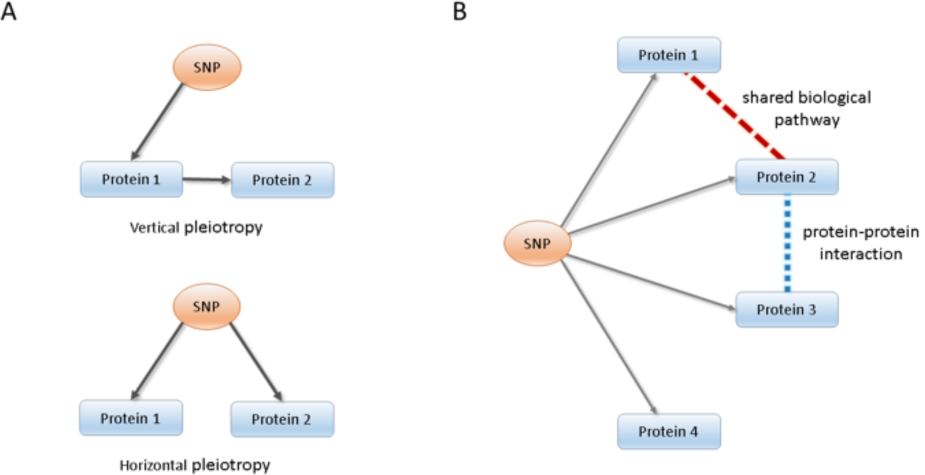
Distinguishing vertical and horizontal pleiotropy using EpiGraphDB. (A) Concept of vertical and horizontal pleiotropy using proteins as an example. We have a valid instrument for MR when a SNP affects proteins in a single path; in contrast, if an instrument is associated with proteins participating in different pathways, it violates the “exclusion restriction criterion” and our instrument is invalid. (B) Integration of SNP-protein associations with pathway information and PPI data to distinguish vertical and horizontal pleiotropy using EpiGraphDB. All four proteins are associated with the same SNP. Protein 1 and protein 2 share the same biological pathway. Protein 2 and 3 are in PPI. Protein 4 shares no links with other proteins. Therefore, the SNP association on protein 1, 2 and 3 are likely to act through vertical pleiotropy, where the SNP association on protein 4 verse other three proteins are likely to be horizontal pleiotropy.

Here, by integrating SNP-protein associations with biological pathway and protein-protein interaction (PPI) information implemented in EpiGraphDB, we have developed an approach to assess potential horizontal pleiotropy. As demonstrated in Figure 2 B, for a SNP associated with a group of proteins, we check the number of biological pathways and PPIs that are shared across this group of proteins. If these proteins are mapped to the same biological pathway and/or a PPI exists between them, then the SNP is more likely to act through vertical pleiotropy and therefore be a valid instrument for MR.

#### 3.1.1 Assessing the pleiotropy of an autoimmune-related variant

In this case study we assessed the pleiotropy of rs12720356, a SNP located in TYK2 gene that is associated with Crohn’s disease and psoriasis (Solovieff *et al.*, 2013), by exploring the relationships between genes (and their products) to which this SNP can be mapped using expression QTL data. We used the GTEx database (The GTEx Consortium *et al.*, 2015) to identify single-tissue eQTL effects, gathering a set of genes whose expression level is associated with rs12720356 in different tissues: FDX1L, ICAM1, ICAM5, KRI1, MRPL4, GRAP2, TMED1, TYK2 and ZGLP6.We then proceeded querying EpiGraphDB to extract pathway and PPI data as described in the methods section.

The results were then converted to a graph that shows two small connected components and a few isolated nodes (Figure 3). ICAM1 shares pathways with ICAM5, the interactions of integrin cell surface and between lymphoid and non-lymphoid cells. Integrin expression has been shown to be altered in psoriasis (Creamer *et al.*, 1995), and integrins also have an important pro-inflammatory role in Crohn’s disease, where they facilitate the movement of leukocytes from the systemic circulation (note that the association is detected in whole blood) to the intestinal mucosa (Park and Jeen, 2018). ICAM1 also participates with TYK2 in the regulation of Interleukin-4 (IL4) and Interleukin-13 (IL3) signalling, important actors that drive a predominantly humorally mediated hypersensitivity response (Sartor, 1994). In terms of PPIs, the above pairs of genes are still connected, and we retrieved a triple formed by ICAM1, RAVER1 and TYK2, and the pair KRI1-MRPL4 that is associated with sun exposure, a well-established beneficial factor for psoriasis and Crohn’s disease (Søyland *et al.*, 2011; Jantchou *et al.*, 2014). However, here the results depict that some single-tissue eQTLs with a strong association, like ZGLP1 and FDX2, remain unconnected in our network. This shows that they potentially work along different molecular pathways, acting in horizontal pleiotropy. It would be important to consider their potential biological role in the outcome phenotypes of any MR analyses using this instrument.

**Fig. 3.**
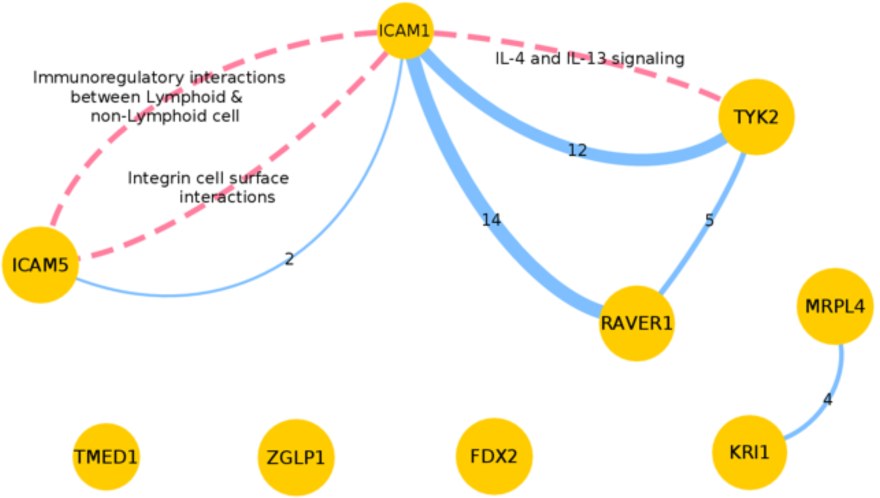
Network diagram with the evidence to assess the pleiotropy of genetic variant rs12720356. The network has one node for each protein regulated by the eQTL rs12720356, and their size is inversely proportional to their P-value (see Supplementary Table 4 for details). Dashed pink edges depict the participation in common biological pathways, and blue edges represent the number of shared PPIs (value indicated).

#### 3.1.2 An exemplar valid instrument

We recently used the same approach to explore potential vertical and horizontal pleiotropy for a number of pleiotropic protein associated SNPs (Zheng, Haberland, *et al.*, 2019). In one example, a specific set of three proteins (IL6ST, ICAM1 and TIMP1) were associated with the same SNP (rs144276707). The pair ICAM1 and TIMP1 were found to participate in two common pathways, and there were 4 shared PPIs among all three proteins. These results supported the hypothesis that rs144276707 is more likely to influence these proteins via the same biological pathway (acting through vertical pleiotropy), strengthening the evidence that this SNP is a valid instrument for MR analysis.

### 3.2 Identification of potential drug targets

Systematic MR of molecular phenotypes such as proteins and levels of transcript expression offers important potential to prioritise drug targets for pharmacological investigation. However, many potential targets are not easily druggable. A parallel problem is that current GWAS of plasma proteins have limited sample size, are not available in many tissues, and only represent a subset of all proteins. A potential way to address these problems is to use protein-protein interaction (PPI) information to identify druggable targets linked to a non-druggable, but robustly causal gene. Their relationship to the causal gene increases our confidence in their potential causal role even if the initial evidence of their causal effect is below our multiple-testing threshold. Here we have developed an approach using PPI data to prioritise potential alternative drug targets. As a proof of principle, we illustrate this approach using IL23R as an example.

#### 3.2.1 Integrating MR evidence with PPI networks for alternative drug targets search

IL23R is a well-established disease-susceptibility gene for inflammatory bowel disease (IBD) (de Lange *et al.*, 2017). The protein-disease association information retrieved from EpiGraphDB suggests that IL23R has a robust causal effect on IBD^7^ (beta = 1.50, P-value = 2.21 × 10^−166^, colocalization probability = 75%) (Zheng, Haberland, *et al.*, 2019). The drug PTG-200, acting as an antagonist of IL23R has just passed Phase I and is in Phase II trials for IBD treatment (Cheng *et al.*, 2019), which aligns well with the genetic/MR evidence implemented in EpiGraphDB. Whilst IL23R is druggable, we illustrate how our approach can identify potential alternative targets using pathway data.

We used PPI information (Orchard *et al.*, 2014; Szklarczyk *et al.*, 2019) and data on druggability (Finan *et al.*, 2017) to identify a set of proteins which are the target of approved drugs and clinical-phase drug candidates and have direct PPI with IL23R. Table 2 shows a subset of this list with strong MR evidence (P-value < 1 × 10^−5^) to IBD^8^.

**Table 2.**
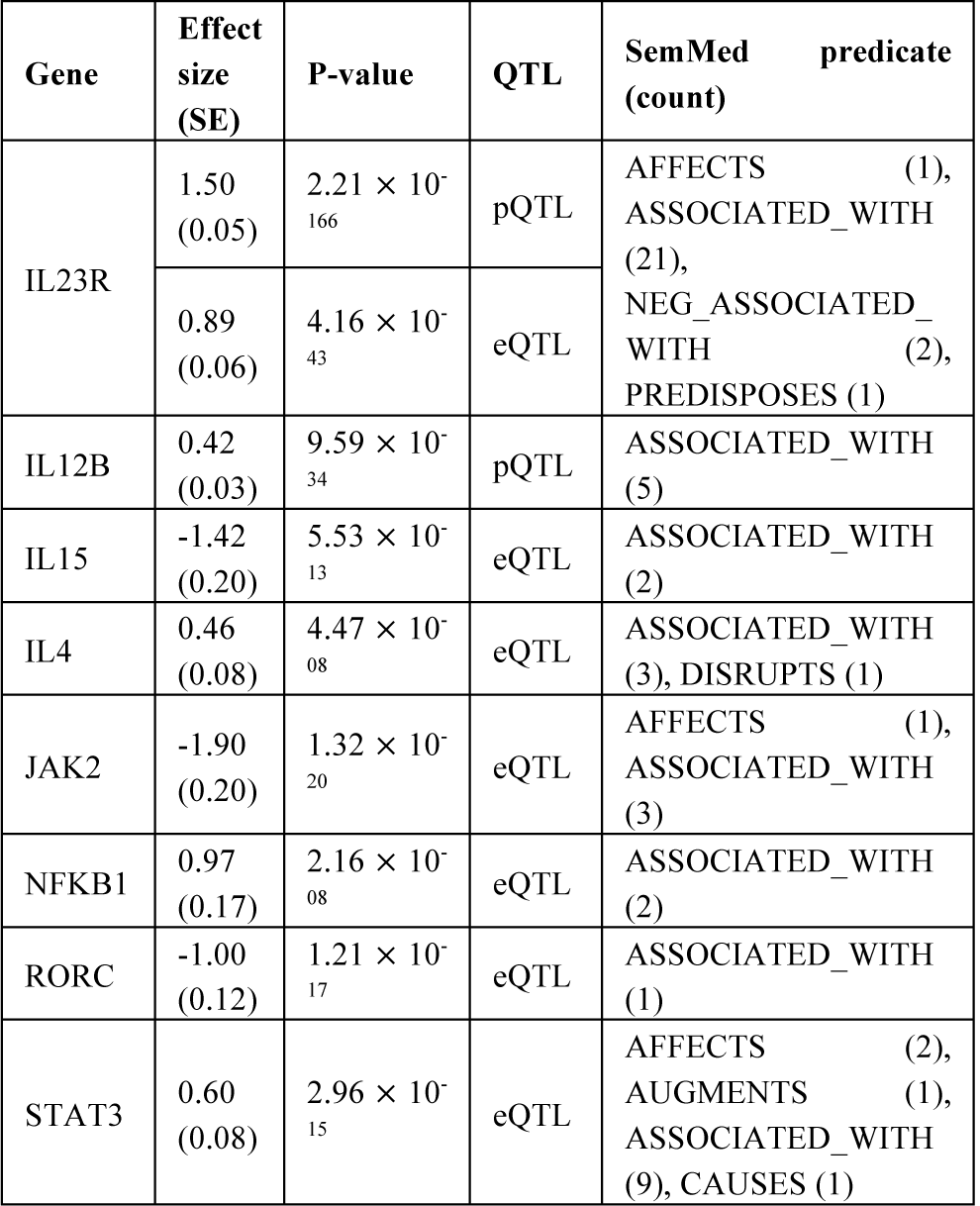
Triangulation of MR and literature evidence on the effects of IL23R and associated genes to IBD. The MR evidence is the QTL MR estimates of IL23R and the associated druggable genes (via direct protein-protein interaction with Tier 1 druggability) to GWAS ieu-a-249: Inflammatory bowel disease. The literature evidence is the SemMed predicates derived by SemMedDB and the numbers of PubMed articles identified to support the predicate mechanism. Here we report the subset of genes that are identified to contain both MR evidence (P-value < 1 × 10-5).

This list of proteins includes IL12B, which is the target protein for an existing drug Ustekinumab, which is currently under Phase 3 and 4 trials for IBD treatment^*9*^. Although there is strong MR evidence for IL12B (beta = 0.42, P-value = 9.59 × 10^−34^), there is little evidence for genetic colocalization^*10*^ (colocalization probability < 1%), which prevents us prioritizing this target based on MR evidence alone. However, the PPI between IL12B and IL23R (which *does* have reliable MR and colocalization results) increases our confidence that IL12B is a valid target.

#### 3.2.2 Using literature evidence for results enrichment and triangulation

A further source of useful evidence is the literature-derived knowledge from SemMedDB (Kilicoglu *et al.*, 2012) available in EpiGraphDB. Integrating this literature evidence with the evidence described above can further enhance confidence in the findings (as well as identify potential alternative drug targets). **Table 2** also reports the gene-to-trait literature evidence regarding IL23R and interacting proteins and IBD, where each entry shows a literature-derived semantic triple (e.g. “IL23R” – “ASSOCIATED_WITH” – “Inflammatory Bowel Diseases”), as well as the study articles from which each triple was extracted. For the list of genes including IL23R and IL12B that were identified with strong MR evidence, we were also able to find abundant literature evidence supporting the genetic causal evidence with derived mechanisms involving predicates such as ASSOCIATED_WITH, AFFECTS and CAUSES.

### 3.3 Triangulating causal estimates with literature evidence

Previously, we have demonstrated that existing literature can be used to derive relationships and mechanisms between defined biomedical traits (Elsworth *et al.*, 2018). By integrating this knowledge with causal estimates in EpiGraphDB, we can triangulate evidence, identifying where these two sources of evidence are in agreement, and where they are not (Lawlor *et al.*, 2017). In this case study we explore the literature connecting traits with pre-defined causal relationships. From here we can summarise the key mechanisms defined in the literature, and also potentially derive novel mechanisms.

#### 3.3.1 Sleep duration and coronary heart disease as an example

Starting with an exposure trait of “Sleep duration”, existing MR data, and connections between traits and diseases in EpiGraphDB, we extracted a set of potentially causally related traits (**Table 3**).

**Table 3.**
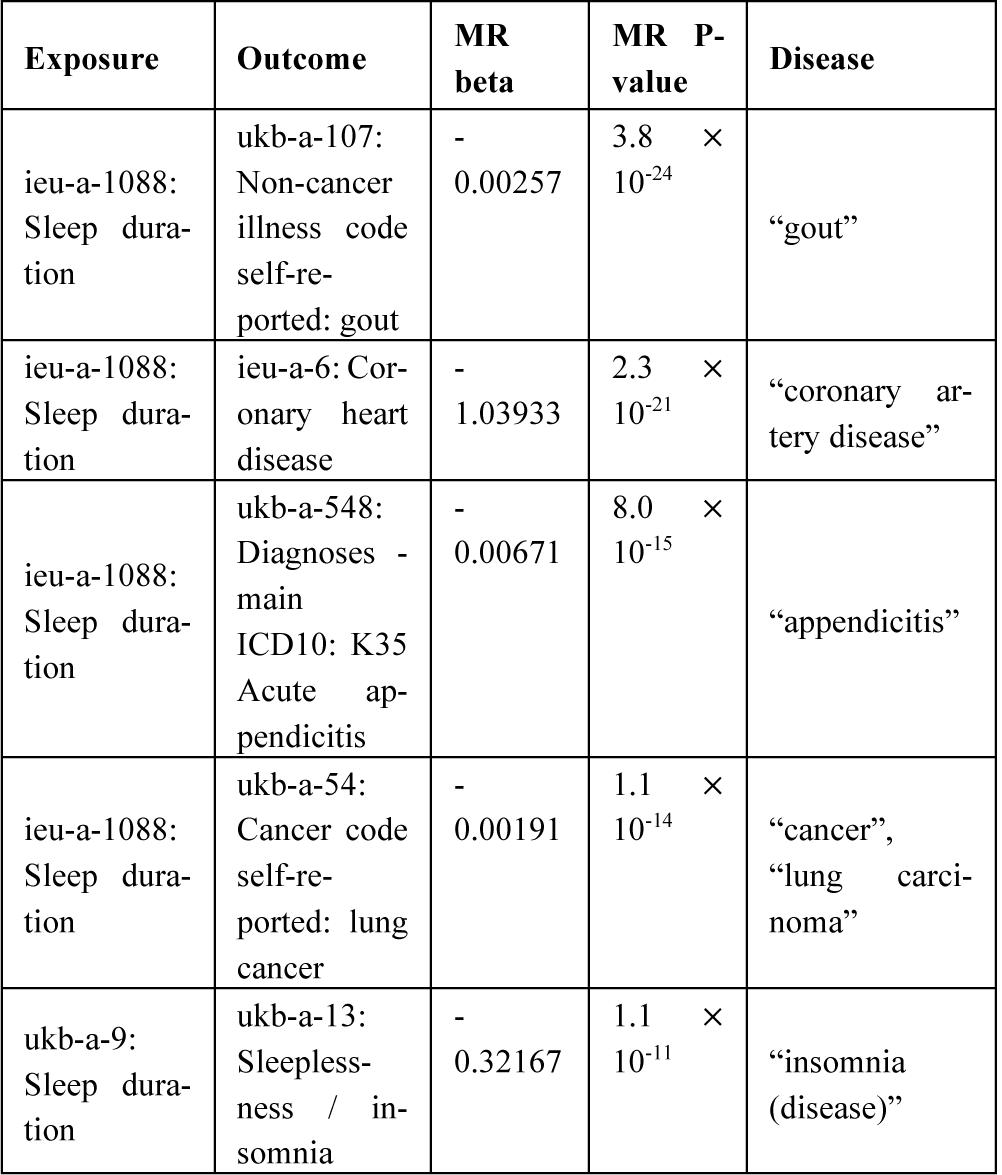
Summary of disease traits identified with causal association to “Sleep duration”. We searched for MR evidence associated with trait “Sleep duration” with P-value to be under 1 × 10^−10^, and map the outcome trait to a disease term via mappings through EFO terms.

Multiple disease entries arise from the mapping between the trait name and EFO terms, each of which maps to a disease term. In this case we treated each as a single relationship and extracted the literature data connecting a pair of traits. For this example, we selected the outcome trait “Coronary heart disease” to explore in more detail the potential mechanisms linking this to sleep duration. To do this we queried EpiGraphDB to extract the semantic triples associated with each trait and searched for overlapping terms, identifying 839 overlapping triples (Supplementary Table 6 reports the top 10 items by enrichment P-value).

We then generated frequency counts for the overlapping terms (Figure 4), which identified many different overlapping terms and types^11^, including 6 proteins (aapp), 2 genes (gngm) and 11/ organic chemicals (orch). Each of these represents a key point in a potential mechanism, connecting the exposure and outcome traits. Terms of particular interest are those with high counts (e.g. Ethanol) as these represent terms with large numbers of supporting publications in the literature. However, in this case, Ethanol may be present in such numbers due to its inclusion in many publications as a co-factor when describing the functionality and efficacy of drugs, highlighting the importance of reviewing a selection of papers underpinning each mechanism.

**Fig. 4.**
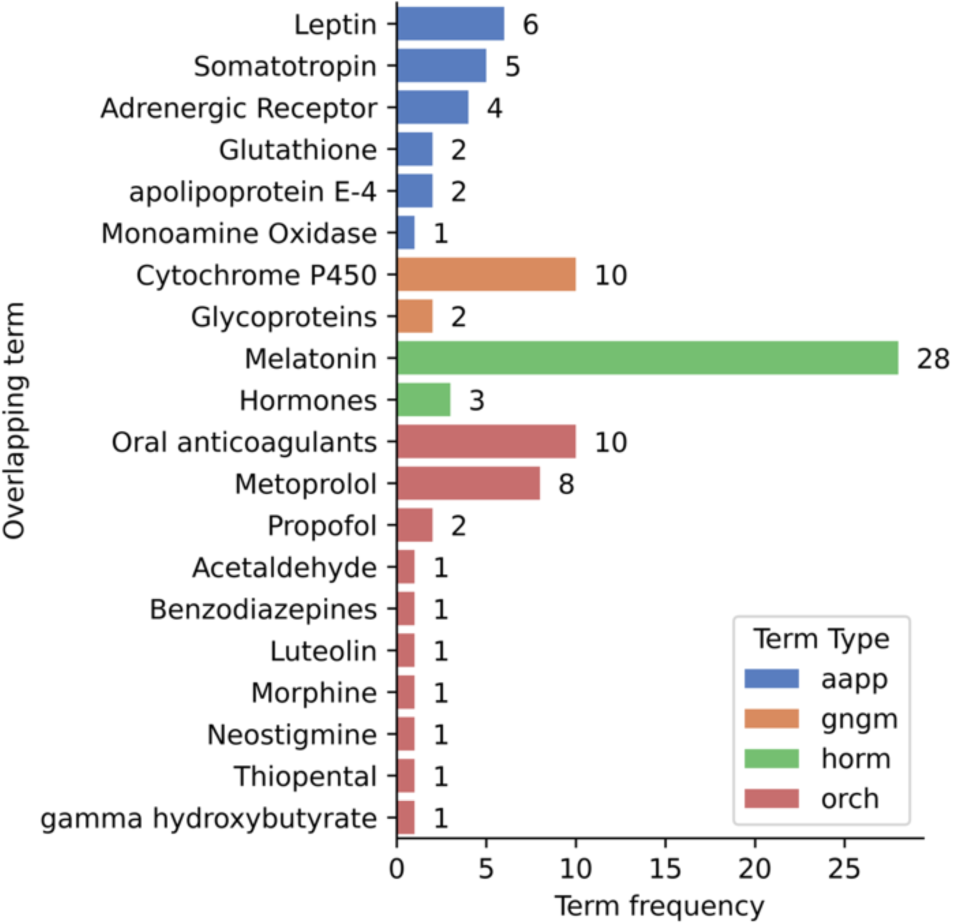
Literature-mined/derived evidence on the intermediates between “Sleep duration” and “Coronary heart disease”. Counts of overlapping SemMed terms grouped by the SemMed term type.

#### 3.3.2 Investigation of one overlapping term in detail

Figure 5 suggests the main route from Sleep Duration to Coronary Heart Disease via the intermediate term Leptin involves only one term on the exposure side (“ghrelin”) and 10 on the outcome side.

**Fig. 5.**
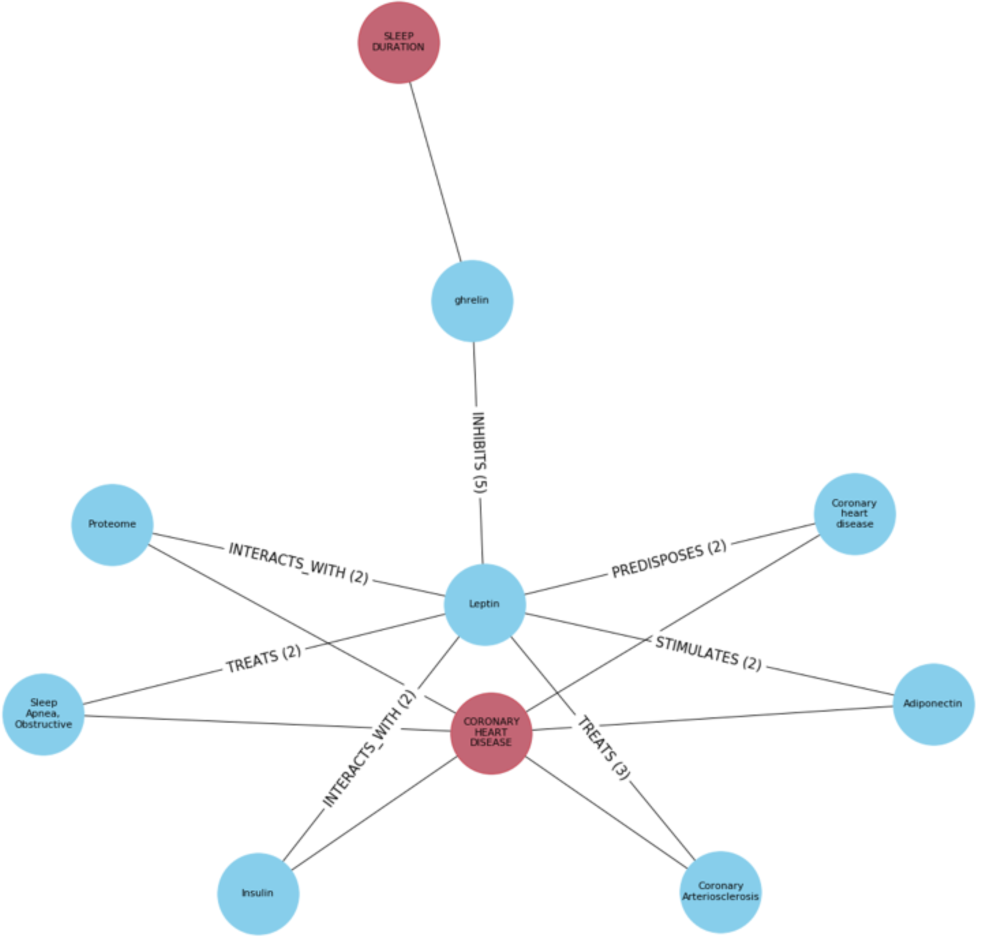
Literature derived mechanisms between “Sleep duration”, “Leptin”, and “Coronary Heart Disease”. Network diagram displaying the literature connections between “Sleep Duration” and “Coronary Heart Disease” through the intermediate term “Leptin”. Predicates connecting two semantic terms and their frequencies are labelled on the edges. Red nodes represent the exposure (SLEEP DURATION) and outcome (CORONARY HEART DISEASE) traits, blue nodes represent intermediate semantic literature nodes.

#### 3.3.3 Check the original publication text

Finally, EpiGraphDB provides a PubMed identifier to enable us to check the validity of these connections in the original text. For example, we found evidence of two statements that “Leptin PREDISPOSES Coronary heart disease”. These were derived from the following two sentences:

> “CONCLUSIONS: Consumption of sugar-sweetened beverages was associated with increased risk of CHD and some adverse changes in lipids, inflammatory factors, and leptin.” (de Koning *et al.*, 2012)
>
> “Leptin, one of the earlier adipocytokines, is known to play a major role in cardiovascular disease and recent observations suggest that leptin is an independent risk factor for coronary heart disease.” (Amasyali *et al.*, 2010)

The contrasting causal interpretation of these two sentences highlights the importance of manual review of the original articles to validate the semantic triples.

## 4 Discussion

EpiGraphDB is a new database and platform for data integration in health data science, with a particular focus on understanding the relationships between risk factors, intermediate phenotypes and disease outcomes described by epidemiological analyses. Whilst we present three specific case studies, we anticipate a much wider array of uses and support this through an open API and R package. It is, however, important to recognise that there are several existing platforms for data integration in the health, biomedical and pharmaceutical domains (Supplementary Table 3).

The Open Targets platform (Koscielny *et al.*, 2017; Carvalho-Silva *et al.*, 2019) (https://www.targetvalidation.org/) integrates a wealth of genomic, phenotypic, ontology and drug target data into a single platform aimed for users in the pharmaceutical industry and research community. Their platform has a well-developed web interface in addition to a comprehensive API and Python package to support use of the API. This open approach has enabled EpiGraphDB to utilise drug/target mappings with Open Targets. However, whilst there is some overlap in this context, the Open Targets platform lacks MR estimates (although it does include genetic association data). Open Targets also includes some literature data, and their LINK platform (https://link.opentargets.io/) extracts semantic relationships from PubMed. However, despite some of the conceptual similarities to EpiGraphDB, their focus is primarily on drug target prioritisation, whilst EpiGraphDB also aims to support evaluation of lifestyle risk factors.

The Hetionet platform (https://het.io/) is a graph database integrating data from more than 29 different databases, which was initially set up to prioritize drugs for repurposing using an innovative approach to predict gene/disease associations (“Project Rephetio”) (Himmelstein and Baranzini, 2015; Himmelstein *et al.*, 2017), but now aims to have a broader remit. The platform is very accessible, with a web application, data downloads in multiple formats and open access to their Neo4j database. The primary focus of the platform is for molecular mechanisms and pharmacologic data while EpiGraphDB additionally encompasses epidemiological relationships (MR causal estimates, genetic correlation, etc) and literature data. However, the open nature of the platform enables users to easily work with Hetionet in parallel with EpiGraphDB.

The Monarch Initiative (Mungall *et al.*, 2017) (https://monarchinitiative.org/) is focused on the integration of genotypic and phenotypic data across species with the aim of identifying related phenotypes and potential animal models of disease. This contrasts with the human-centric epidemiological focus of EpiGraphDB. The Monarch Initiative platform has an open source approach to software development and offers web interfaces powered by an open API. In common with Hetionet and EpiGraphDB, the platform uses the Neo4j database. Users can easily integrate data from the Monarch Initiative with EpiGraphDB given their open design principles.

Wikidata (https://wikidata.org) is a general knowledge base which contains an array of biomedical data sources that have recently been reported (Waagmeester *et al.*, 2020). In contrast to curated knowledge bases such as EpiGraphDB, Wikidata is developed through community driven efforts and bot automation, and incorporates extensive knowledge across a wide array of fields, including (but not limited to) a range of biomedical entities, with duplication and redundancy of entities inevitable. This much broader approach also distinguishes Wikidata from specialist platforms such as EpiGraphDB, which is focused on epidemiological and biomedical knowledge. In common with other platforms listed above, the open design of this platform supports cross-platform data integration.

Various other platforms (Gaspar *et al.*, 2018; Coker *et al.*, 2019; Abbot *et al.*, 2020) exist with some conceptual overlaps with EpiGraphDB (Supplementary Table 3). These represent a range of different types of data based on molecular and genetic interactions and drug targets. However, in contrast to the platforms described above these platforms don’t appear to have accessible API or software packages. Although several are open access and available to the wider community, the lack of programmatic interoperability limits their scope.

As with all similar platforms, EpiGraphDB is constrained by the available data and subject to any errors or quality issues that exist in original sources. However, by integrating data from a range of sources (e.g. STRING, IntAct and Reactome for interactions between proteins) we ensure the user can evaluate consistency between data sources.

## 5 Conclusions

The EpiGraphDB platform provides an integrated data resource to support data mining and interpretation of the relationships between disease risk factors, intervention targets and disease outcomes. We present three illustrative case studies that demonstrate the functionality and utility of the platform, but it is important to note that much more extensive capabilities are available and will continue to expand as the platform is developed further. We aim to support open science by making the data freely accessible, both programmatically and through a web interface, and by providing open source code and exemplar Jupyter notebooks.

## Author contribution

YL, BE, PE, VH and TRG wrote and edited the manuscript. YL, BE, TRG developed the platform architecture, and YL, BE, TRG, VH developed the platform components. BE developed the data integration pipeline, and BE, YL, PE, VH, GH, ML, JZ were responsible for the data generation, acquisition and curation. YL, PE, VH, BE, JZ developed the case studies. TRG conceptualised and supervised the project. All authors contributed to the review of the paper and provided comments.

## Footnotes

1 Documentation on the endpoint can be found at https://docs.epigraphdb.org/api/api-endpoints/-post-cypher.

2 https://epigraphdb.org/cypher.

3 We refer to a type of biomedical entity as a *meta node* (e.g. `(Gwas)` in Cypher notation) and a type of association as a *meta relationship* (e.g. `[MR]`), whereas a specific entity is referred to as a *node* (e.g. `(Gwas {id: “ieu-a-2”, trait: “Body mass index”})`) and a specific association as a *relationship* (e.g. `(Gwas {trait: “Body mass index”})-[MR {beta, se, pval}]->(Gwas {trait: “Coronary heart disease”})`).

4 Further details on the inhouse results by EpiGraphDB members are available from Appendix 2 in the supplementary materials.

5 Information and metrics are based on latest version of EpiGraphDB platform (version 0.3.0, 21 April 2020).

6 Details on the list of pleiotropic genes are reported in Supplementary Table 4.

7 https://epigraphdb.org/pqtl/IL23R

8 Supplementary Table 5 reports the full list of identified proteins with druggability information.

9 Drug trial information available via Open Targets https://www.target-validation.org/evidence/ENSG00000113302/EFO_0000540?view=sec:known_drug

10 http://epigraphdb.org/pqtl/IL12B

11 https://mmtx.nlm.nih.gov/MMTx/semanticTypes.shtml

