## supplementary materials for "EpiGraphDB: A database and data mining platform for health data science"

### Appendix 1 Biomedical entities and associations in EpiGraphDB

Biomedical entities are represented as nodes in EpiGraphDB and associations (including causal evidence as well as mappings and similarities of entities) are represented as relationships. For example, a GWAS (ieu-a-2) on body mass index from OpenGWAS is represented as `(Gwas {id: “ieu-a-2”, trait: “Body mass index”})` in the Neo4j Cypher notation in EpiGraphDB. The MR evidence between the GWAS of body mass index and coronary heart disease is represented as `(Gwas {id: “ieu-a-2”, trait: “Body mass index”})-[MR {beta, se, p}]->(Gwas {id: “ieu-a-9”, “Coronary heart disease”})` with effect size and other coefficients / categories represented as properties of the relationship. In addition, we refer to a type of entities such as GWAS as a meta node `(Gwas)`, and a type of associations such as MR as a meta relationship `[MR]`.

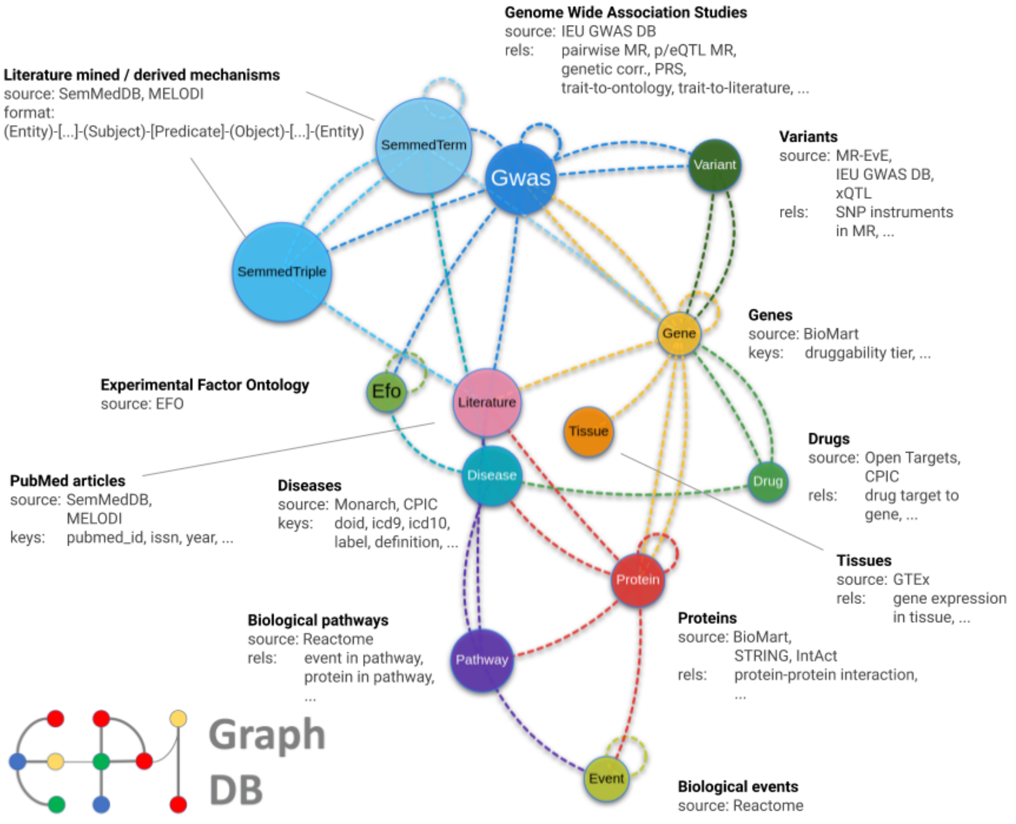

**Supplementary Fig. 1. Integrated Epidemiological Evidence in EpiGraphDB**. Each node represents a meta node which is a type of biomedical entity and each link represents a meta relationship (e.g. causal associations or mappings) between two meta nodes. A live version of the schema can be accessed at https://epigraphdb.org/about - schema.

**Supplementary Table 1. EpiGraphDB meta nodes**

| Meta node | Entity count | Source |
|---|---|---|
| `(Disease)` | 21,829 | Monarch (Mungall *et al.*, 2017) |
| `(Drug)` | 2,455 | CPIC (Relling and Klein, 2011), Open Targets (Carvalho-Silva *et al.*, 2019), (Finan *et al.*, 2017) |
| `(Efo)` | 25,390 | EFO (Malone *et al.*, 2010) |
| `(Event)` | 11,868 | Reactome (Jassal *et al.*, 2019) |
| `(Gene)` | 59,171 | BioMart (Smedley *et al.*, 2009) |
| `(Gwas)` | 31,773 | OpenGWAS (Elsworth, Lyon, *et al.*, 2020) |
| `(Literature)` | 29,137,785 | SemMedDB (Kilicoglu *et al.*, 2012), MELODI (Elsworth *et al.*, 2018) |
| `(Pathway)` | 2,180 | Reactome (Jassal *et al.*, 2019) |
| `(Protein)` | 21,543 | BioMart (Smedley *et al.*, 2009) |
| `(SemmedTerm)` | 105,047 | SemMedDB (Kilicoglu *et al.*, 2012), MELODI (Elsworth *et al.*, 2018) |
| `(SemmedTriple)` | 3,428,531 | SemMedDB (Kilicoglu *et al.*, 2012), MELODI (Elsworth *et al.*, 2018) |
| `(Variant)` | 88,125 | MR-EvE (Hemani *et al.*, 2017), OpenGWAS (Elsworth, Lyon, *et al.*, 2020), xQTL (Zheng *et al.*, 2019) |
| `(Tissue)` | 53 | GTEx (Lonsdale *et al.*, 2013) |

* Information and metrics are based on the latest version of EpiGraphDB platform (version 0.3.0, 21 April 2020).

**Supplementary Table 2. EpiGraphDB meta relationships**

| Meta relationship | Source | Target | Entity Count | Source |
|---|---|---|---|---|
| *[MONDO_MAP_EFO]* | Disease | Efo | 2,822 | Monarch (Mungall *et al.*, 2017) |
| *[MONDO_MAP_UMLS]* | Disease | SemmedTerm | 3,417 | Monarch (Mungall *et al.*, 2017) |
| *[OPENTARGETS_DRUG_TO_DISEASE]* | Drug | Disease | 2,486 | Open Targets (Carvalho-Silva *et al.*, 2019) |
| *[OPENTARGETS_DRUG_TO_TARGET]* | Drug | Gene | 6,024 | Open Targets (Carvalho-Silva *et al.*, 2019) |
| *[CPIC]* | Drug | Gene | 355 | CPIC (Relling and Klein, 2011) |
| *[EFO_CHILD_OF]* | Efo | Efo | 43,154 | EFO (Malone *et al.*, 2010) |
| *[PRECEDING_EVENT]* | Event | Event | 10,418 | Reactome (Jassal *et al.*, 2019) |
| *[XQTL_SINGLE_SNP_MR_GENE_GWAS]* | Gene | Gwas | 8,703,863 | xQTL (Zheng *et al.*, 2019) |
| *[XQTL_MULTI_SNP_MR]* | Gene | Gwas | 3,098,049 | xQTL (Zheng *et al.*, 2019) |

| | | | | |
|---|---|---|---|---|
| *[GENE_TO_LITERATURE]* | Gene | Literature | 771 | SemMedDB (Kilicoglu *et al.*, 2012), MELODI (Elsworth *et al.*, 2018) |
| *[INTACT_INTERACTS_WITH_GENE_PROTEIN]* | Gene | Protein | 1,451 | IntAct (Orchard *et al.*, 2014) |
| *[GENE_TO_PROTEIN]* | Gene | Protein | 20,762 | Reactome (Jassal *et al.*, 2019) |
| *[EXPRESSED_IN]* | Gene | Tissue | 861,552 | Reactome (Jassal *et al.*, 2019) |
| *[GWAS_NLP_EFO]* | Gwas | Efo | 6,936 | Vectology (Elsworth, Liu, *et al.*, 2020) |
| *[MR]* | Gwas | Gwas | 583,619 | MR-EvE (Hemani *et al.*, 2017) |
| *[BN_GEN_COR]* | Gwas | Gwas | 904,832 | Neale Lab (Abbot *et al.*, 2020) |
| *[OBS_COR]* | Gwas | Gwas | 17,932 | EpiGraphDB inhouse |
| *[GWAS_NLP]* | Gwas | Gwas | 30,838,964 | Vectology (Elsworth, Liu, *et al.*, 2020) |
| *[PRS]* | Gwas | Gwas | 132,703 | PRS Atlas (Richardson *et al.*, 2019) |

| | | | | |
|---|---|---|---|---|
| *[GWAS_TO_LIT]* | Gwas | Literature | 19,079,468 | SemMedDB (Kilicoglu *et al.*, 2012), MELODI (Elsworth *et al.*, 2018) |
| *[METAMAP_LITE]* | Gwas | SemmedTerm | 2,081 | MetaMap (Demner-Fushman *et al.*, 2017) |
| *[GWAS_SEM]* | Gwas | SemmedTriple | 9,075,020 | SemMedDB (Kilicoglu *et al.*, 2012), MELODI (Elsworth *et al.*, 2018) |
| *[GWAS_TO_VARIANT]* | Gwas | Variant | 26,521 | MR-EvE (Hemani *et al.*, 2017) |
| *[TOPHITS]* | Gwas | Variant | 122,730 | OpenGWAS (Elsworth, Lyon, *et al.*, 2020) |
| *[PATHWAY_TO_DISEASE]* | Pathway | Disease | 541 | Reactome (Jassal *et al.*, 2019) |
| *[EVENT_IN_PATHWAY]* | Pathway | Event | 12,488 | Reactome (Jassal *et al.*, 2019) |
| *[PATHWAY_TO_LITERATURE]* | Pathway | Literature | 8,952 | SemMedDB (Kilicoglu *et al.*, 2012) |
| *[PROTEIN_TO_DISEASE]* | Protein | Disease | 626 | Reactome (Jassal *et al.*, 2019) |
| *[PROTEIN_IN_EVENT]* | Protein | Event | 13,484 | Reactome (Jassal *et al.*, 2019) |

| | | | | |
|---|---|---|---|---|
| *[PROTEIN_TO_LITERATURE]* | Protein | Literature | 107,315 | SemMedDB (Kilicoglu *et al.*, 2012) |
| *[PROTEIN_IN_PATHWAY]* | Protein | Pathway | 9,955 | Reactome (Jassal *et al.*, 2019) |
| *[INTACT_INTERACTS_WITH_PROTEIN_PROTEIN]* | Protein | Protein | 187,426 | IntAct (Orchard *et al.*, 2014) |
| *[STRING_INTERACT_WITH]* | Protein | Protein | 390,222 | STRING (Szklarczyk *et al.*, 2019) |
| *[INTACT_NOT_INTERACTS_WITH]* | Protein | Protein | 699 | IntAct (Orchard *et al.*, 2014) |
| *[SEM_GENE]* | SemmedTerm | Gene | 41,706 | SemMedDB (Kilicoglu *et al.*, 2012) |
| *[SEM_PREDICATE]* | SemmedTerm | SemmedTerm | 3,428,531 | SemMedDB (Kilicoglu *et al.*, 2012) |
| *[SEM_TO_LIT]* | SemmedTriple | Literature | 6,127,985 | SemMedDB (Kilicoglu *et al.*, 2012) |
| *[SEM_SUB]* | SemmedTriple | SemmedTerm | 3,428,531 | SemMedDB (Kilicoglu *et al.*, 2012) |
| *[SEM_OBJ]* | SemmedTriple | SemmedTerm | 3,428,531 | SemMedDB (Kilicoglu *et al.*, 2012) |

| | | | | |
|---|---|---|---|---|
| *[XQTL_SINGLE_SNP_MR_SNP_GENE]* | Variant | Gene | 41,564 | xQTL<br>(Zheng *et al.*, 2019) |
| *[VARIANT_TO_GENE]* | Variant | Gene | 59,157 | MR-EvE<br>(Hemani *et al.*, 2017) |

* Information and metrics are based on the latest version of EpiGraphDB platform (version 0.3.0, 21 April 2020).

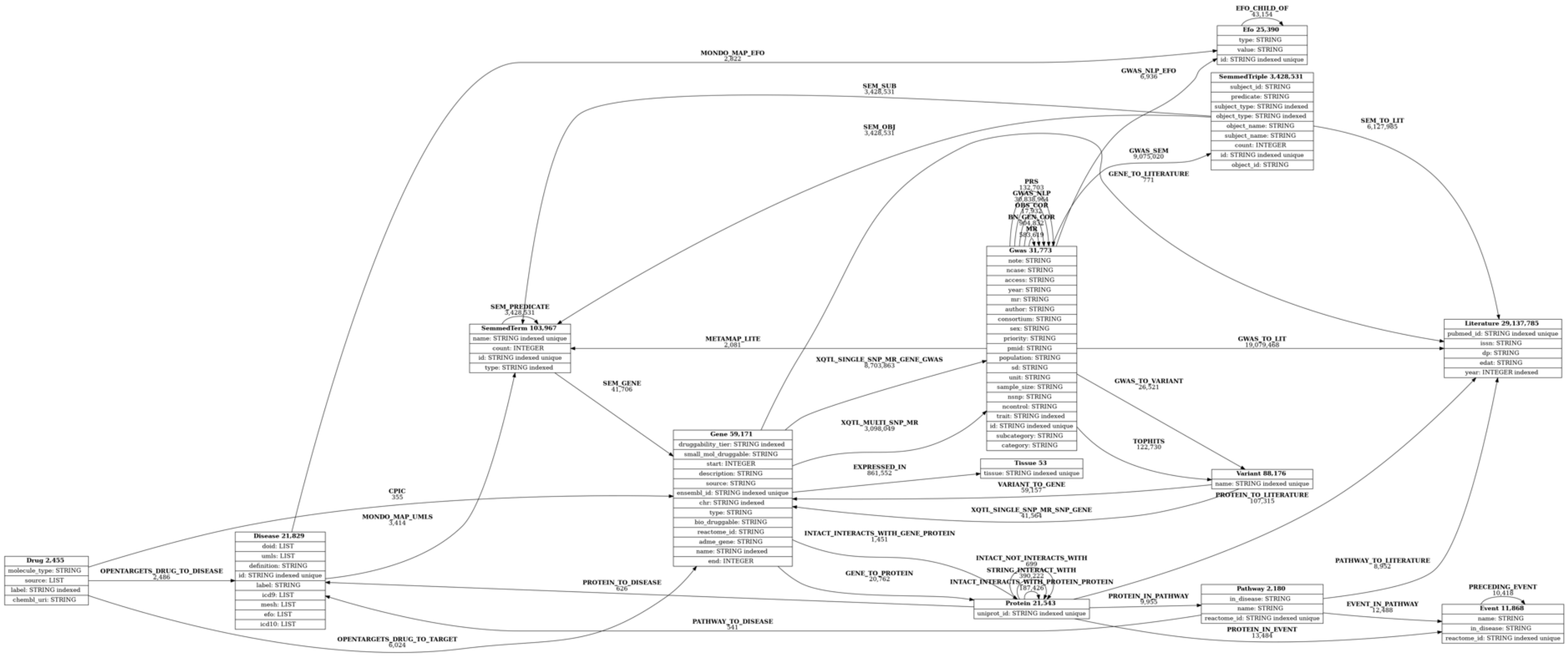

***Supplementary Fig. 2. EpiGraphDB Graph schema***. *Each box represents a meta node which is a class of biomedical entity and each link represents a meta relationship between two meta nodes. A live version of the schema can be accessed from the EpiGraphDB API https://api.epigraphdb.org/meta/schema?graphviz=true&plot=true.*

#### Appendix 2 Data integration

To facilitate a flexible and dynamic development environment, we developed a data integration and graph construction method composed of multiple integration methods wrapped up in a SnakeMake pipeline (Köster and Rahmann, 2018). For large data sets, e.g. millions of rows, we used the Neo4j import tool https://neo4j.com/docs/operations-manual/current/tutorial/import-tool/ which will ingest 100's of millions of rows in minutes. Data ingested in this way need to be in simple pre-processed text files, avoiding duplicates and conflicting entries. After this bulk ingest, the remaining data are added using the LOAD CSV https://neo4j.com/docs/cypher-manual/current/clauses/load-csv/ method. This is slower than the import tool, however, it provides flexibility during the import, e.g. checking for existing entries, modifying entries on insert, etc. Whilst collating and creating the data ready for insert takes many hours, building the graph takes ~15 minutes, providing an agile modification and rebuild method.

The following gives a brief description of each data set, the source of the raw data and the order in which they were inserted into the graph.

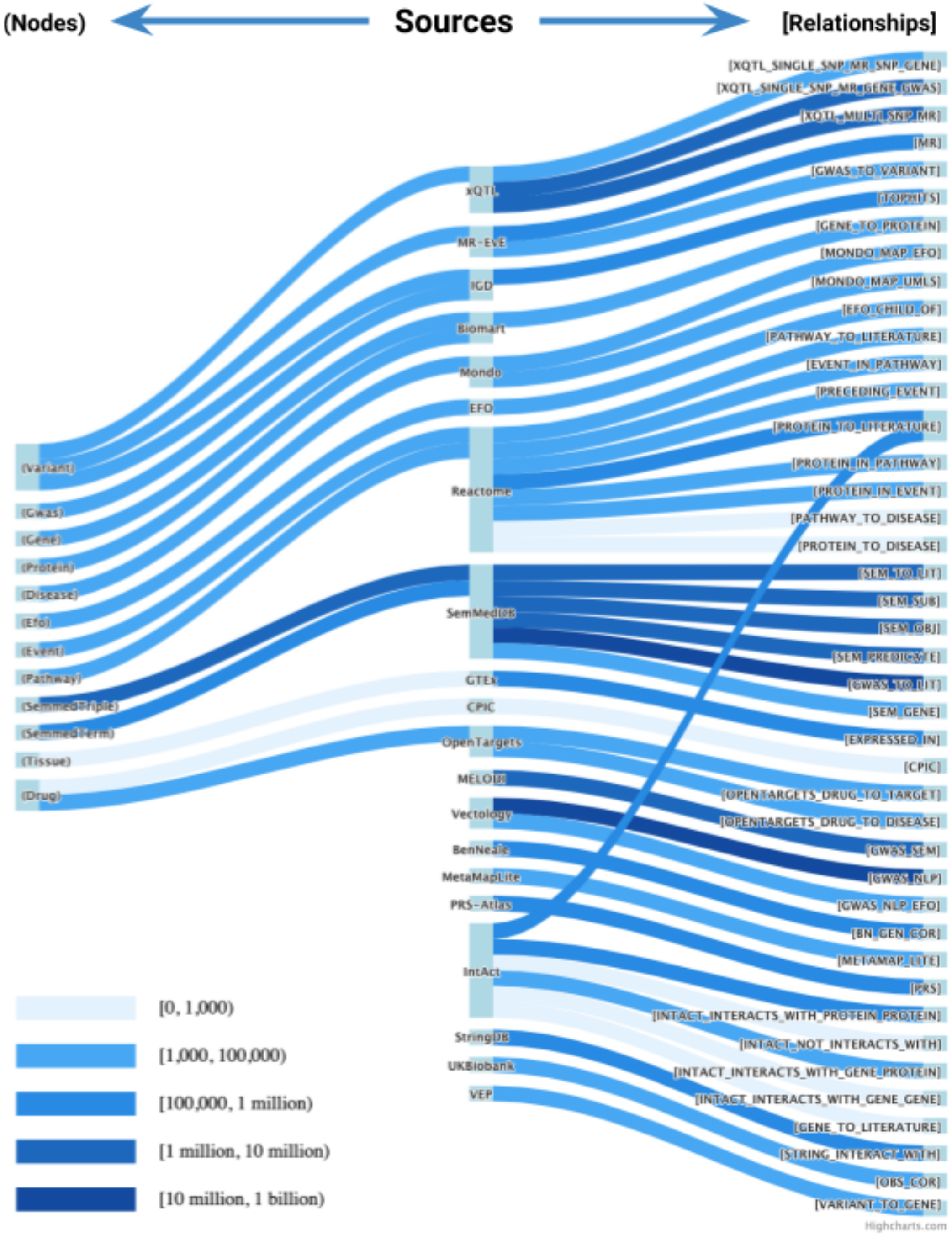

***Supplementary Fig. 3. Integration of source data into EpiGraphDB.*** *Entities in the middle represent the sources of data, and entities on the left and right represent meta nodes and meta relationships in EpiGraphDB. A live version of the data integration flow diagram can be accessed at https://epigraphdb.org/about - data-integration.*

#### 1 Genetic variants

Source: OpenGWAS (Elsworth, Lyon, *et al.*, 2020), MR-EvE (Hemani *et al.*, 2017) and xQTL (Zheng *et al.*, 2019)

Multiple sources of genetic variant data were combined into a single set of unique variants. We resisted adding a complete set of data, e.g. dbSNP, instead opting to reduce the size of the graph, and only include variants with links to other data. This included top hits for each GWAS (variants with LD $R^2 < 0.001$ within a 10Mb window and with P-value $< 5 \times 10^{-8}$), variants derived from MR-EvE and the xQTL data. This unique list of variants was annotated with potential gene effects using a local instance of Ensembl Variant Effect Predictor (VEP) v37 (McLaren *et al.*, 2016).

**2 Genome wide association studies**

Source: OpenGWAS (Elsworth, Lyon, *et al.*, 2020)

The IEU OpenGWAS platform contains GWAS summary data for over 11,000 full and 20,000 partial data sets. Each full data set represents a measured variable (phenotype) relating to a specific area of human health and each partial data set represents the set of quantitative trait loci (QTL) for a particular molecular trait (e.g. gene expression, methylation or metabolite levels). Molecular QTL data are typically not genome-wide, focusing instead on cis genomic regions and strongly associated trans-QTL. For each GWAS we incorporated the meta data for and the subset of genetic variants most associated with that trait into EpiGraphDB. These data were extracted via the OpenGWAS API (http://gwas-api.mrcieu.ac.uk/docs/).

**3 Genes and Proteins**

Source: BioMart (Smedley *et al.*, 2009)

We used Biomart build 37 via the biomart python client https://pypi.org/project/biomart/ to create our set of genes and proteins. For simplicity and to avoid issues with conflicting IDs, we restricted our gene search to gene names and Ensembl IDs, and for proteins we used UniProt IDs only.

**4 Diseases**

Source: Mondo Disease Ontology (Mondo) (Mungall *et al.*, 2017)

Representing disease knowledge is problematic as there are numerous ontologies and sources of information. We chose the Mondo Disease Ontology (Mondo) as it has the explicit aim to harmonize disease definitions across different ontologies, and appears to be the most complete such resource, providing links to many other ontologies.

**5 Observational Correlation**

Source: UK Biobank (UK Biobank, 2014)

We used PHESANT (Millard *et al.*, 2018) to create a modified version of variables from the UK Biobank https://www.ukbiobank.ac.uk/. Using a basic spearman rank correlation, we produced pairwise correlation coefficients for all UK Biobank phenotypes for which we also have GWAS results from OpenGWAS (https://gwas.mrcieu.ac.uk/datasets/). These pairwise correlation coefficients create observational correlation relationships between the `Gwas` nodes.

**6 Mendelian Randomization**

Source: MR-EvE (Hemani *et al.*, 2017)

Using a mixture of experts model, MR-EvE estimated pairwise causal relationships using Mendelian randomization (MR) between many of the traits for which we have GWAS results integrated from OpenGWAS. These data are used to create MR relationships between the `Gwas` nodes.

SemMedDB is a collection of semantic relations, created from the titles and abstracts of PubMed. Each record is a triple (subject-predicate-object) with the subject and object mapping to either the UMLS Metathesaurus (National Library of Medicine, 2009) or Entrez gene ID (National Center for Biotechnology Information, 2020). We included unique semantic triple data from the PREDICATION table of SemMedDB (version semmedVER40_R) with terms and predicates matching specific criteria (Elsworth, 2020).

**8 GWAS Literature**

Source: OpenGWAS (Elsworth, Lyon, *et al.*, 2020), SemMedDB (Kilicoglu *et al.*, 2012)

MELODI (Elsworth *et al.*, 2018) uses literature annotations, e.g. SemMedDB to derive connections between two search terms, e.g. an exposure and outcome. Using a modified version of this approach (Elsworth, 2020) which restricts the search space to particular subjects, objects and predicates, we created enriched literature objects for each GWAS trait, using the GWAS trait text. These were then be used to identify overlapping terms between two traits. The derivation of these literature connections for a pair of traits is as follows:

- For each trait a PubMed search was performed (using the trait name).

- The PubMed IDs returned by this search were used to retrieve matching triples of data (subject-predicate-object) from a local instance of SemMedDB (v40).
- Each triple was counted and compared to the background count to produce an enrichment P-value.
- Enriched, overlapping triples (object from exposure, subject from outcome) were returned and form the literature relationship between pairs of traits.

**9 GWAS semantic similarity**

Source: OpenGWAS (Elsworth, Lyon, *et al.*, 2020)

Many of the GWAS in OpenGWAS are of a similar nature. For example, a search of "weight" returns 24 records https://gwas.mrcieu.ac.uk/datasets/?trait__icontains=weight, and there are many similar trait names that don't contain the word "weight". We developed a platform to help address the issue of understanding the sematic similarity between biomedical variables which uses sentence embedding (Elsworth, Liu, *et al.*, 2020). This can be used to search a set of variables using a model of knowledge extracted from the biomedical literature in place of simple text matching, and can also be used to create distance/similarity scores between two pieces of text. In this case, we compared the sentence embedding vectors derived from the GWAS traits to create similarity scores.

**10 Pathways**

Source: Reactome (Jassal *et al.*, 2019)

For biological pathway data we utilised a subset of data available from Reactome. To avoid overloading our graph we extracted a reduced set of information from the Reactome graph, focusing on the connections between pathways and events, literature, disease and proteins.

**11 Drugs**

Source: Open Targets (Carvalho-Silva *et al.*, 2019)

Open Targets is a platform focused on drug targets and discovery, containing data from many sources and provides straightforward programmatic access to specific components. Using their API, we extracted data related to drug, trial phase and target gene.

**12 Drug efficacy**

Source: Clinical Pharmacogenetics Implementation Consortium (CPIC) (Relling and Klein, 2011)

CPIC is an international consortium interested in facilitating the clinical implementation of pharmacogenetic tests. They provide a freely available database of peer-reviewed, evidence-based gene/drug clinical practice guidelines, including systematic grading of evidence. We retrieved from this resource data about drug efficacy.

**13 Tissue specific gene expression**

Source: Genotype Tissue Expression (GTEx) project (The GTEx Consortium *et al.*, 2015)

Tissue-specific gene expression levels were obtained from GTEx. This public resource contains samples from 54 non-diseased tissue sites across nearly 1,000 individuals.

**14 Protein-protein interactions (1)**

Source: IntAct (Orchard *et al.*, 2014)

IntAct is a freely available, open source database of molecular interaction data derived from literature curation and direct user submissions. We distilled the subset of protein-protein interactions (PPI) to incorporate it in our database.

**15 Protein-protein interactions (2)**

Source: StringDB (Szklarczyk *et al.*, 2019)

StringDB is another database that contains known and predicted protein-protein interactions, either direct or functional (indirect). Data comes from computational predictions of PPI in different organisms. We selected the direct PPIs for homo sapiens with a confidence probability greater than 0.7.

**16 Druggable genes**

Source: The druggable genome study (Finan *et al.*, 2017)

Finan et al. (2017) presented an approach to validate drug targets using genomic information. Data from genome-wide association studies were used to connect complex disease- and biomarker-associated loci, meaning that associations of variants in genes that encode a target mimic the effect of modifying pharmacologically these targets. From there we extracted a set of genes encoding druggable human proteins.

**17 Genetic correlations**

Source: Neale Lab UK Biobank genetic correlation study (Abbot *et al.*, 2020)

In October 2019 the Ben Neale Lab released genetic correlation data for over 4,000 GWAS. We incorporated the correlation data that mapped to the GWAS trait names in EpiGraphDB.

**18 GWAS trait to UMLS**

Source: OpenGWAS (Elsworth, Lyon, *et al.*, 2020)

MetaMap Lite https://metamap.nlm.nih.gov/MetaMapLite.shtml (Demner-Fushman *et al.*, 2017) was used to create a relationship between the GWAS traits names and SemMedDB terms (based on the UMLS metathesaurus).

**19 Polygenic Risk Scores**

Source: PRS Atlas (Richardson *et al.*, 2019)

Richardson et al. (2019) conducted a systematic analysis on the association between 162 polygenic risk scores for different phenotypes (PRS; derived from published GWAS in OpenGWAS) and 551 traits from UK Biobank. We integrated the associations provided by this study (http://mrcieu.mrsoftware.org/PRS_atlas/) by mapping the PRS and UK Biobank to the EpiGraphDB GWAS traits, creating new relationships representing the published PRS associations.

**20 pQTL and eQTL MR**

Source: xQTL study (Zheng *et al.*, 2019)

Zheng et al. (2019) conducted systematic MR and colocalization analyses of 1,740 plasma proteins (pQTLs) and 16,058 blood transcripts (eQTLs) on 576 phenotypes in Europeans on the drug target prioritization for complex diseases. We integrated these results in EpiGraphDB for both single SNP MR results and multi SNP MR results (IVW and Egger MR methods).

**21 Experimental Factor Ontology**

Source: EFO (Malone *et al.*, 2010)

In addition to a disease ontology we required an established ontology that covered a broad range of biomedical variables. We selected EFO https://www.ebi.ac.uk/efo/ as a well-established and comprehensive ontology. Each class and parent-child relationship were downloaded via the EBI SPARQL endpoint (https://www.ebi.ac.uk/rdf/services/sparql) on January 28th 2020.

To create links between GWAS and EFO, we utilised the Vectology platform (Elsworth, Liu, *et al.*, 2020). This used sentence embedding methods to create vectors of each GWAS trait and EFO term, then performed a simple distance measure to identify closest matches. This was chosen in preference to alternative methods, such as Zooma (https://www.ebi.ac.uk/spot/zooma/) and OnToma (https://ontoma.readthedocs.io/en/stable) as we experienced superior results using embedding methods.

### Appendix 3 Comparison of data integration platforms

**Supplementary Table 3. Comparison of data integration platforms**

| **Platform** | **URL** | **Focus** | **WebApp** | **Accessible API** | **R/Python package** | **Open Access** | **Database architecture** |
|---|---|---|---|---|---|---|---|
| EpiGraphDB | https://epigraphdb.org/ | Lifestyle and drug target prioritization, identification of potential molecular mechanisms and pleiotropy | Y | Y | R | Y | Neo4j |
| Open Targets | https://www.opentargets.org/ | Integration of phenotype, genomic, literature and drug data to support identification of new targets for drug discovery. | Y | Y | Python | Y | ClickHouse DBMS |
| Hetionet | https://het.io/ | Biomedical relationships to support hypothesis generation and mechanistic insights (including drug repurposing) | Y | Y | Python | Y | Neo4j |
| Monarch initiative | https://monarchinitiative.org/ | Data integration and analytics integrating phenotype and genotype across species to evaluate phenotype-based similarity | Y | Y | Various | Y | Neo4j |
| Wikidata | https://www.wikidata.org | Comprehensive knowledge base, including biomedical entities | Y | Y | 3rd party packages | Y | Blazegraph |

| | | | | | | | |
|---|---|---|---|---|---|---|---|
| rg | https://ukbb-rg.hail.is/ | UK Biobank genetic correlation browser and dataset | Y | N | N | Y | TSV files |
| canSAR Black | https://cansarblack.icr.ac.uk/ | Integration of multi-disciplinary data to support machine learning for drug discovery in cancer | Y | N | N | Y | Unclear |
| Navigome | https://phenviz.navigome.com/ | Visualization of phenotype cluster, genetic correlations and pathways. Discovery of characteristic genes/pathways for phenotypes | Y | N | N | Y | Unclear |

### Appendix 4: Query EpiGraphDB

Here we provide a simple demonstration of querying EpiGraphDB using the standard API endpoints, and the R package, and lastly Cypher. We query for MR evidence between phenotypes which “Sleep duration” is the exposure phenotype. For examples of programmatic usage of EpiGraphDB please visit the Jupyter notebooks https://github.com/MRCIEU/epigraphdb.

#### Query via EpiGraphDB API

The EpiGraphDB API https://api.epigraphdb.org provides a self-documenting interface (Swagger) where users can interactively adjust the parameters and view the returned data on the spot. In addition to the query results, the response from the API returns the associated Cypher query that is used to query from EpiGraphDB Graph. We provide detailed documentation on the usage, parameters and scenarios for the API endpoints on the project documentation site https://docs.epigraphdb.org/. Here we show how to query the API service using the Requests library in Python.

```
from pprint import pprint
import requests

api_url = "https://api.epigraphdb.org"
endpoint = "/mr"
response = requests.get(
    api_url + endpoint,
    params={
      "exposure": "Sleep duration",
    }
)
response.raise_for_status()
result = response.json()

pprint(result)
# {'metadata': {'empty_results': False,
#               'query': 'MATCH (exposure:Gwas)-[mr:MR]->(outcome:Gwas) WHERE '
#                        'exposure.trait = "Sleep duration" AND mr.pval < 1e-05 '
#                        'RETURN exposure {.id, .trait}, outcome {.id, .trait}, '
#                        'mr {.b, .se, .pval, .method, .selection, .moescore} '
#                        'ORDER BY mr.pval ;',
#               'total_seconds': 0.013717},
#  'results': [{'exposure': {'id': 'ieu-a-1088', 'trait': 'Sleep duration'},
#               'mr': {'b': -0.010066916234791279,
#                      'method': 'FE IVW',
#                      'moescore': 1.0,
#                      'pval': 0.0,
#                      'se': 0.00013109980500303211,
#                      'selection': 'DF'},
#               'outcome': {'id': 'ukb-a-460',
#                           'trait': 'Vitamin and mineral supplements: Vitamin '
```

```
#                             'B'}},
#              {'exposure': {'id': 'ieu-a-1088', 'trait': 'Sleep duration'},
#               'mr': {'b': -0.18189099431037903,
#                      'method': 'FE IVW',
#                      'moescore': 1.0,
#                      'pval': 0.0,
#                      'se': 0.011486822739243507,
#                      'selection': 'DF'},
#               'outcome': {'id': 'ieu-a-118', 'trait': 'Neuroticism'}},
#              {'exposure': {'id': 'ieu-a-1088', 'trait': 'Sleep duration'},
#               'mr': {'b': 2.3293509483337402,
#                      'method': 'FE IVW',
```

Query via `epigraphdb` R package

The equivalent query using the R package is[1]:

```
library("epigraphdb")
result = mr(exposure_trait="Sleep duration")

print(result)
# # A tibble: 104 x 12
# exposure_id exposure_name outcome_id outcome_name estimate      se     p
# <chr>       <chr>         <chr>      <chr>           <dbl>   <dbl> <dbl>
# 1 1088        Sleep durati… UKB-a:460  Vitamin and…  -0.0101 1.31e-4     0
# 2 1088        Sleep durati… 581        X-12093         0.103  6.22e-4     0
# 3 1088        Sleep durati… 668        Laurylcarni…  -0.0750 1.27e-3     0
# 4 1088        Sleep durati… 386        Phenylaceta…   0.0360 2.90e-4     0
# 5 1088        Sleep durati… 315        Pyruvate       0.0782 1.18e-3     0
# 6 1088        Sleep durati… 432        Aspartylphe…   0.133  5.15e-4     0
# 7 1088        Sleep durati… 454        X-10810       -0.0364 8.13e-5     0
# 8 1088        Sleep durati… 1158       Stearic aci…   0.450  6.35e-3     0
# 9 1088        Sleep durati… 1065       Inspection …  -0.498  4.85e-4     0
# 10 1088        Sleep durati… 1073       Copper         0.250  6.39e-3     0
# # … with 94 more rows, and 5 more variables: ci_upp <dbl>, ci_low <dbl>,
# #   selection <chr>, method <chr>, moescore <dbl>
```

[1] The epigraphdb R package is hosted on Github and can be installed using devtools in R as `devtools::install_github("MRCIEU/epigraphdb-r").` For eash of use by default the epigraphdb R package will convert the result JSON object into a tabular data frame, which can be switched off to return the same data structure of the EpiGraphDB API. Please refer to the package documentation https://mrcieu.github.io/epigraphdb-r for its usage.

#### Query using Cypher

EpiGraphDB supports querying the Neo4j graph database using the Cypher[2] query language from API (via POST /cypher). Here we show how the above request can be achieved using the `GET /cypher` endpoint.

##### Python requests

```
from pprint import pprint
import requests

api_url = https://api.epigraphdb.org
endpoint = "/cypher"
response = requests.post(
    api_url + endpoint,
    json={
      "query": 'MATCH (exposure:Gwas)-[mr:MR]->(outcome:Gwas) WHERE exposure.trait
= "Sleep duration" AND mr.pval < 1e-05 RETURN exposure {.id, .trait}, outcome {.id
, .trait}, mr {.b, .se, .pval, .method, .selection, .moescore} ORDER BY mr.pval ;'
,
    }
)
response.raise_for_status()
result = response.json()

pprint(result)
# {'metadata': {'empty_results': False,
#               'query': 'MATCH (exposure:Gwas)-[mr:MR]->(outcome:Gwas) WHERE '
#                        'exposure.trait = "Sleep duration" AND mr.pval < 1e-05 '
#                        'RETURN exposure {.id, .trait}, outcome {.id, .trait}, '
#                        'mr {.b, .se, .pval, .method, .selection, .moescore} '
#                        'ORDER BY mr.pval ;',
#               'total_seconds': 0.284867},
#  'results': [{'exposure': {'id': 'ieu-a-1088', 'trait': 'Sleep duration'},
#               'mr': {'b': -0.010066916234791279,
#                      'method': 'FE IVW',
#                      'moescore': 1.0,
#                      'pval': 0.0,
#                      'se': 0.00013109980500030321,
#                      'selection': 'DF'},
#               'outcome': {'id': 'ukb-a-460',
#                           'trait': 'Vitamin and mineral supplements: Vitamin '
#                                    'B'}},
#              {'exposure': {'id': 'ieu-a-1088', 'trait': 'Sleep duration'},
#               'mr': {'b': -0.18189099431037903,
#                      'method': 'FE IVW',
#                      'moescore': 1.0,
#                      'pval': 0.0,
#                      'se': 0.011486822739243507,
#                      'selection': 'DF'},
```

[2] For detailed tutorials on Cypher please refer to Neo4J's documentation https://neo4j.com/developer/cypher-basics-i/.

```
#                'outcome': {'id': 'ieu-a-118', 'trait': 'Neuroticism'}},
#               {'exposure': {'id': 'ieu-a-1088', 'trait': 'Sleep duration'},
#                'mr': {'b': 2.3293509483337402,
#                       'method': 'FE IVW',
```

R package

```
library("epigraphdb")

endpoint = "/cypher"
params = list(query='MATCH (exposure:Gwas)-[mr:MR]->(outcome:Gwas) WHERE
exposure.trait = "Sleep duration" AND mr.pval < 1e-05 RETURN exposure {.id,
.trait}, outcome {.id, .trait}, mr {.b, .se, .pval, .method, .selection,
.moescore} ORDER BY mr.pval')

result = query_epigraphdb(
    route=endpoint,
    params=params,
    mode="table",
    method="POST"
)
print(result)
# # A tibble: 104 x 12
# exposure_id exposure_name outcome_id outcome_name estimate      se     p
# <chr>       <chr>         <chr>      <chr>           <dbl>   <dbl> <dbl>
# 1 1088        Sleep durati… UKB-a:460  Vitamin and…  -0.0101 1.31e-4     0
# 2 1088        Sleep durati… 581        X-12093        0.103  6.22e-4     0
# 3 1088        Sleep durati… 668        Laurylcarni…  -0.0750 1.27e-3     0
# 4 1088        Sleep durati… 386        Phenylaceta…   0.0360 2.90e-4     0
# 5 1088        Sleep durati… 315        Pyruvate       0.0782 1.18e-3     0
# 6 1088        Sleep durati… 432        Aspartylphe…   0.133  5.15e-4     0
# 7 1088        Sleep durati… 454        X-10810       -0.0364 8.13e-5     0
# 8 1088        Sleep durati… 1158       Stearic aci…   0.450  6.35e-3     0
# 9 1088        Sleep durati… 1065       Inspection …  -0.498  4.85e-4     0
# 10 1088        Sleep durati… 1073       Copper         0.250  6.39e-3     0
# # … with 94 more rows, and 5 more variables: ci_upp <dbl>, ci_low <dbl>,
# #   selection <chr>, method <chr>, moescore <dbl>
```

### Appendix 5: Case studies

#### Case study 1: Distinguishing vertical and horizontal pleiotropy for SNP-protein associations

We retrieved from GTEx database the single-tissue eQTL effects for the SNP of interest, rs12720356, identifying 9 genes whose expression level is associated with it. We then queried EpiGraphDB to extract pathway and PPI data and conduct the analyses as follows:

- First, we mapped each of the identified genes to their coding proteins using the `POST /mappings/gene-to-protein` endpoint in EpiGraphDB, retrieving one mapped protein for each gene in our database.
- We then used the `POST /protein/in-pathway` endpoint to identify the Reactome pathways[3] each protein participates in, either if these proteins are involved in the pathway as a single entity or as part of a complex.
- For each pair of proteins we looked for shared pathways and we converted these results into a graph, using proteins as nodes and pathways as edges. The numbers of connected components in the graph were counted.
- For each pair of proteins we retrieved shared PPIs from STRING (Szklarczyk *et al.*, 2019) data using the `POST /protein/ppi/pairwise` endpoint in EpiGraphDB. Note that we considered either direct PPI or an interaction with one mediator protein (i.e. a protein that interacts with both proteins in our pair). The number of PPIs per pair of proteins was counted, and it was used to build a graph, where we looked at the number of connected components.

[3] Pathway ontology is structured in a hierarchical way, but here we considered only the bottom layer of the hierarchy.

The analyses presented here can be replicated using the code available at https://github.com/MRCIEU/epigraphdb/blob/master/paper-case-studies/case-1-pleiotropy.ipynb.

**Supplementary Table 4. List of genes associated with the SNP of interest rs12720356.** Significant single tissue eQTLs for the variant, from GTEx[1]. For each protein, we queried EpiGraphDB[2] to obtain the number of pathways that they are involved in and total number of direct protein-protein interactions.

| HGNC name | P-value[3] | Pathways | PPIs | Tissue |
|---|---|---|---|---|
| FDX1L | $2.90 \times 10^{-11}$ | 0 | 13 | thyroid, nerve, brain, testis, adipose, artery |
| ICAM1 | $1.00 \times 10^{-04}$ | 2 | 147 | whole blood |
| ICAM5 | $7.60 \times 10^{-07}$ | 1 | 3 | whole blood |
| KRI1 | $9.90 \times 10^{-06}$ | 0 | 130 | skin[a] |
| MRPL4 | $6.80 \times 10^{-09}$ | 0 | 192 | nerve, fibroblasts, skin[a,b], artery, adipose |
| GRAP2 | $5.40 \times 10^{-06}$ | 0 | 71 | whole blood |
| TMED1 | $4.80 \times 10^{-05}$ | 0 | 4 | whole blood |
| TYK2 | $3.00 \times 10^{-06}$ | 1 | 107 | whole blood, adrenal gland |
| ZGLP1 | $1.10 \times 10^{-15}$ | 0 | 1 | thyroid, brain[c], nerve, testis, skin[b], prostate, oesophagus, pituitary |

[a]sun exposed; [b]not sun exposed; [c]includes multiple tissues

[1]GTEx V8, retrieved 21 April 2020. Ensembl gene identifiers were converted to gene name using EpiGraphDB.

[2]EpiGraphDB version 0.3.0, retrieved 21 April 2020.

[3]The smallest value for each gene is reported.

#### Case study 2: Identification of potential drug targets

We used the `GET /gene/druggability/ppi` endpoint of EpiGraphDB to extract proteins with known PPIs with IL23R from IntAct (Orchard *et al.*, 2014) and STRING (Szklarczyk *et al.*, 2019), with additional annotations on druggability (Finan *et al.*, 2017). In our analyses we focused on "Tier 1" targets, for which the gene is the target gene for approved drugs and clinical-phase drug candidates.

We extracted MR evidence for the effects of these genes on IBD using the `GET /xqtl/single-snp-mr` endpoint in EpiGraphDB, which provides causal estimates based on both pQTL and eQTL data.

Finally, we used the `GET /gene/literature` endpoint to extract information from the literature that links the potential alternative targets with IBD.

The analyses presented here can be replicated using the code available at https://github.com/MRCIEU/epigraphdb/blob/master/paper-case-studies/case-2-alt-drug-target.ipynb.

**Supplementary Table 5. Identification of alternative druggable genes to IL23R in PPI network.** We queried EpiGraphDB on genes that directly interact with IL23R in the protein-protein interaction network from IntAct (Orchard *et al.*, 2014) and STRING (Szklarczyk *et al.*, 2019) where we limit to genes that contain druggability information from the druggable genome study (Finan *et al.*, 2017).

| HGNC name | UniProt ID | Druggability tier |
|---|---|---|
| CSF2 | P04141 | Tier 1 |
| IFNA1 | P01562 | Tier 1 |
| IFNG | P01579 | Tier 1 |
| IL10 | P22301 | Tier 1 |
| IL12B | P29460 | Tier 1 |
| IL12RB1 | P42701 | Tier 1 |
| IL13 | P35225 | Tier 1 |
| IL15 | P40933 | Tier 1 |

| | | |
|---|---|---|
| IL17A | Q16552 | Tier 1 |
| IL17F | Q96PD4 | Tier 1 |
| IL2 | P60568 | Tier 1 |
| IL22 | Q9GZX6 | Tier 1 |
| IL23A | Q9NPF7 | Tier 1 |
| IL4 | P05112 | Tier 1 |
| IL5 | P05113 | Tier 1 |
| IL6 | P05231 | Tier 1 |
| IL9 | P15248 | Tier 1 |
| JAK1 | P23458 | Tier 1 |
| JAK2 | O60674 | Tier 1 |
| NFKB1 | P19838 | Tier 1 |
| PIK3CA | P42336 | Tier 1 |
| RORC | P51449 | Tier 1 |
| STAT3 | P40763 | Tier 1 |
| TSLP | Q969D9 | Tier 1 |
| TYK2 | P29597 | Tier 1 |
| CCR6 | P51684 | Tier 2 |
| NFKBIA | P25963 | Tier 2 |
| NOD2 | Q9HC29 | Tier 2 |
| PIK3R1 | P27986 | Tier 2 |
| RELA | Q04206 | Tier 2 |
| STAT1 | P42224 | Tier 2 |
| STAT5A | P42229 | Tier 2 |
| STAT6 | P42226 | Tier 2 |
| CSF3 | P09919 | Tier 3A |

| | | |
|---|---|---|
| ERAP1 | Q9NZ08 | Tier 3A |
| IL12A | P29459 | Tier 3A |
| IL17D | Q8TAD2 | Tier 3A |
| IL19 | Q9UHD0 | Tier 3A |
| IL21 | Q9HBE4 | Tier 3A |
| IL24 | Q13007 | Tier 3A |
| IL7 | P13232 | Tier 3A |
| CCRL2 | O00421 | Tier 3B |

#### Case study 3: Triangulating causal estimates with literature evidence

MELODI uses data from PubMed and SemMedDB (Kilicoglu *et al.*, 2012) to identify enriched overlapping terms between two sets of literature. We utilized the MELODI Presto version (Elsworth, 2020) (which offers significant performance improvements and an API) to create enriched literature terms for all traits (with some defined filtering steps).

We used the /mr endpoint to identify traits for which sleep duration is a potential causal exposure. We then used the `GET /ontology/gwas-efo-disease` endpoint to identify the subset of these traits which are diseases (according to Experimental Factor Ontology). We then selected one specific disease outcome (coronary heart disease) to illustrate the literature mining approach.

The `GET /literature/gwas/pairwise` endpoint was used to identify potential intermediates using the approach we have previously described (Elsworth *et al.*, 2018). The `GET /literature/gwas` endpoint was used in conjunction with MELODI Presto to extract the text from original articles which underpin the semantic triples used to infer intermediate mechanisms.

The analyses presented here can be replicated using the code available at https://github.com/MRCIEU/epigraphdb/blob/master/paper-case-studies/case-3-literature-triangulation.ipynb.

**Supplementary Table 6. Top 10 overlapping SemMedDB triples for "Sleep duration" and "Coronary heart disease".** We searched for the overlapping SemMed triples for "Sleep duration" and "Coronary heart disease". Here we showed the top 10 entries ordered by enrichment p-value.

| s1.subject_name | s1.predicate | s1.object_name / s2.subject_name | s2.predicate | s2.object_name |
|---|---|---|---|---|
| ghrelin | INHIBITS | Leptin | ASSOCIATED_WITH | Coronary heart disease |
| ghrelin | INHIBITS | Leptin | TREATS | Coronary Arteriosclerosis |
| ghrelin | INHIBITS | Leptin | ASSOCIATED_WITH | Coronary Arteriosclerosis |
| ghrelin | INHIBITS | Leptin | PREDISPOSES | Coronary heart disease |
| ghrelin | INHIBITS | Leptin | INTERACTS_WITH | Proteome |
| ghrelin | INHIBITS | Leptin | COEXISTS_WITH | ID1 |

| | | | | |
|---|---|---|---|---|
| ghrelin | INHIBITS | Leptin | COEXISTS_WITH | Hydrocortisone |
| ghrelin | INHIBITS | Leptin | ASSOCIATED_WITH | Cardiovascular Diseases |
| ghrelin | INHIBITS | Leptin | ASSOCIATED_WITH | Cerebrovascular accident |
| ghrelin | INHIBITS | Leptin | TREATS | Sleep Apnea, Obstructive |
